# The Ser83, Arg85, Tyr88, Asn124, Lys192 of C-terminal Lipid-associated membrane hemagglutinin affecting *Mycoplasma synoviae* agglutination of erythrocyte

**DOI:** 10.64898/2026.04.08.717210

**Authors:** Duo-Duo Si, Shi-Jun Bao, Lei Guo, Xiu-Hong Chen, Feng-Qin Wong, Xiao-Xiao He, Qing Wang, Yuan Shi, Sheng-Hu He, Ji-Dong Li

## Abstract

*Mycoplasma synoviae* is an avian pathogen that causes respiratory disease and synovitis, and its hemagglutinin plays a critical role in host cell adhesion. However, the key residues and structural mechanisms underlying hemagglutination remain unclear. In this study, domain analysis of the hemagglutinin family of *Mycoplasma synoviae* revealed that it contains long-chain and short-chain types, among which LAM HA (VY93_RS01465) was selected as the bait protein due to its complete C-terminal conserved domain. Through yeast two-hybrid screening, 18 host proteins interacting with LAM HA were identified. Furthermore, five key amino acid residues S83, R85, Y88, N124, and K192 were found to mediate hemagglutination activity. Deletion of these residues reduced the hemagglutination titer of LAM HA under acidic conditions. Secondary structure analysis showed that the deletion mutation decreased the α-helix content while increasing the proportions of β-sheet and random coil. Molecular dynamics simulations revealed that the mutant exhibited generally higher root mean square deviation and root mean square fluctuation values than the wild-type under different pH conditions, with a marked decrease in structural stability particularly at pH 5.0 and 6.0. These findings indicate that LAM HA, as a critical adhesin, exerts its hemagglutination function dependent on specific key residues and pH-sensitive conformational stability.

**IMPORTANCE:** *Mycoplasma synoviae* (*M. synoviae*) causes significant economic losses to the poultry industry worldwide. Lipid-related membrane protein hemagglutinin (LAM HA) is a surface adhesin essential for host cell attachment, but its precise amino acid residues and structural features have not been defined. In this study, five key residues (S83, R85, Y88, N124, and K192) were identified as critical for LAM HA-mediated hemagglutination activity. Deletion of these residues altered the secondary structure composition, reduced conformational stability under acidic pH conditions, and decreased hemagglutination activity. These findings reveal a previously unknown structure-function relationship of *M. synoviae* LAM HA, demonstrating that its hemagglutination activity depends on specific residues and pH-sensitive structural integrity. This provides new insights into the molecular mechanisms of *M. synoviae* adhesion and offers potential targets for the development of novel intervention strategies against avian mycoplasmosis.

## Introduction

*Mycoplasma synoviae* is an important avian pathogen that causes arthritis and air sac inflammation in chickens and turkeys, resulting in considerable economic losses to the poultry industry worldwide. Due to the lack of a cell wall, several mycoplasma membrane components can directly interact with host cells. Adhesion is the initial step in pathogenic infection and colonization, and cell adhesion-related proteins may therefore play a critical role in pathogenesis.

Hemagglutinin, as an adhesion protein, has been extensively studied in viral pathogenesis. Different hemagglutinin subtypes exhibit significant variability in binding to host cell receptors, directly affecting viral pathogenicity and transmissibility. For instance, hemagglutinins of the H5N1 subtype can switch from avian to human receptors through specific amino acid mutations, a key event for cross-species transmission ^(1, 2)^. The hemagglutinin cleavage site also serves as a critical determinant of viral virulence; alterations in its amino acid sequence-such as those observed in the H5N2 subtype—can markedly enhance pathogenicity and transmission ^(2, 3)^. In *M. synoviae*, hemagglutinin is an abundant surface-exposed lipoprotein generated via post-translational cleavage of the vlhA gene ^(4)^. The 5’ vlhA region encodes a proline-rich repeat at the amino terminus of MSPB, which exhibits high strain-to-strain polymorphism and induces NO secretion in chicken macrophages, as well as IL-6 and IL-1β in chicken monocytes. MSPA possesses hemagglutinin activity and primarily mediates mycoplasma binding to erythrocytes ^(5)^. Recombination between the 5′ vlhA gene and pseudogenes in the genome drives antigenic variation within the C-terminal two-thirds of MSPB and MSPA, thereby altering the domains involved in erythrocyte binding ^(6)^. A conserved motif, P-X-(BCAA)-X-F-X-(BCAA)-X-A-K-X-G, has been identified as a sialic acid–binding motif that may mediate hemagglutinin-dependent attachment to host cells ^[7]^. The C-terminal region of MSPA elicits the highest titers of antibodies capable of inhibiting hemagglutination, hemadsorption, and metabolism ^(7)^. Additionally, the membrane surface of *M. synoviae* is enriched in LAMP, which play a critical role in adhesion and invasion. Three *M. synoviae* hemagglutinins have been identified as LAMP and are predicted to function as adhesins, facilitating both host cell adhesion and erythrocyte agglutination.

Protein function is closely linked to electrostatic interactions, and solution pH regulates key biological processes—including protein folding, ligand binding, enzymatic catalysis, and transmembrane transport—by modulating the protonation states of ionizable residues ^(8)^. Molecular dynamics (MD) simulations can serve as a complement to traditional experiments by capturing transient and cryptic states, thereby bridging static structures with biological function ^(9, 10)^. In this study, we characterize the structural features of three LAM HA domains and screen for host cell bait proteins. By identifying proteins that interact with LAM HA, we aim to elucidate the key amino acid residues involved in erythrocyte agglutination and investigate the influence of pH on this function. The stability of LAM HA under varying pH conditions is assessed through MD simulations. In-depth exploration of these key functional sites will provide a critical theoretical foundation for monitoring *M. synoviae* infection and for the development of vaccines and targeted therapeutics.

## Result

### Conservative domain analysis

The hemagglutinin sequence of 13 *M. synoviae* strains was extracted from the genome data published by NCBI. The results showed that there were two proteins with different lengths in the coding region of the hemagglutinin gene. One peptide chain was composed of 298-385 (average 343) amino acid residues, and the other peptide chain was composed of 142-180 (average 155) amino acid residues. In each specific *M. synoviae* strain, the number of ORFs in the long-chain group (15∼22) was about twice that in the short-chain group (6∼11). Analysis of two types of peptide chains, there is no sequence conserved site between the long-chain group and the short-chain group, but there are certain conserved structural sites in the long-chain group or the short-chain group between the same strain or different strains, which may be related to its function (Fig 1A). Hemagglutinin on the surface of cell membrane is an important adhesin of *M. synoviae*. In a previous study, three hemagglutinins were counted among 200 highly abundantly expressed LAMPs with NCBI numbers VY93_RS04120, VY93_RS04075, and VY93_RS01465, respectively ^(11)^. All three LAM HAs belong to the long chain group, and occur differential expression when *M. synoviae* infects host cells (Fig 1C). LAM HA (VY93_RS01465) containing all the C-terminal conserved structural domains and used as a bait protein for screening of reciprocal host proteins (Fig 1B).

**Fig 1.**
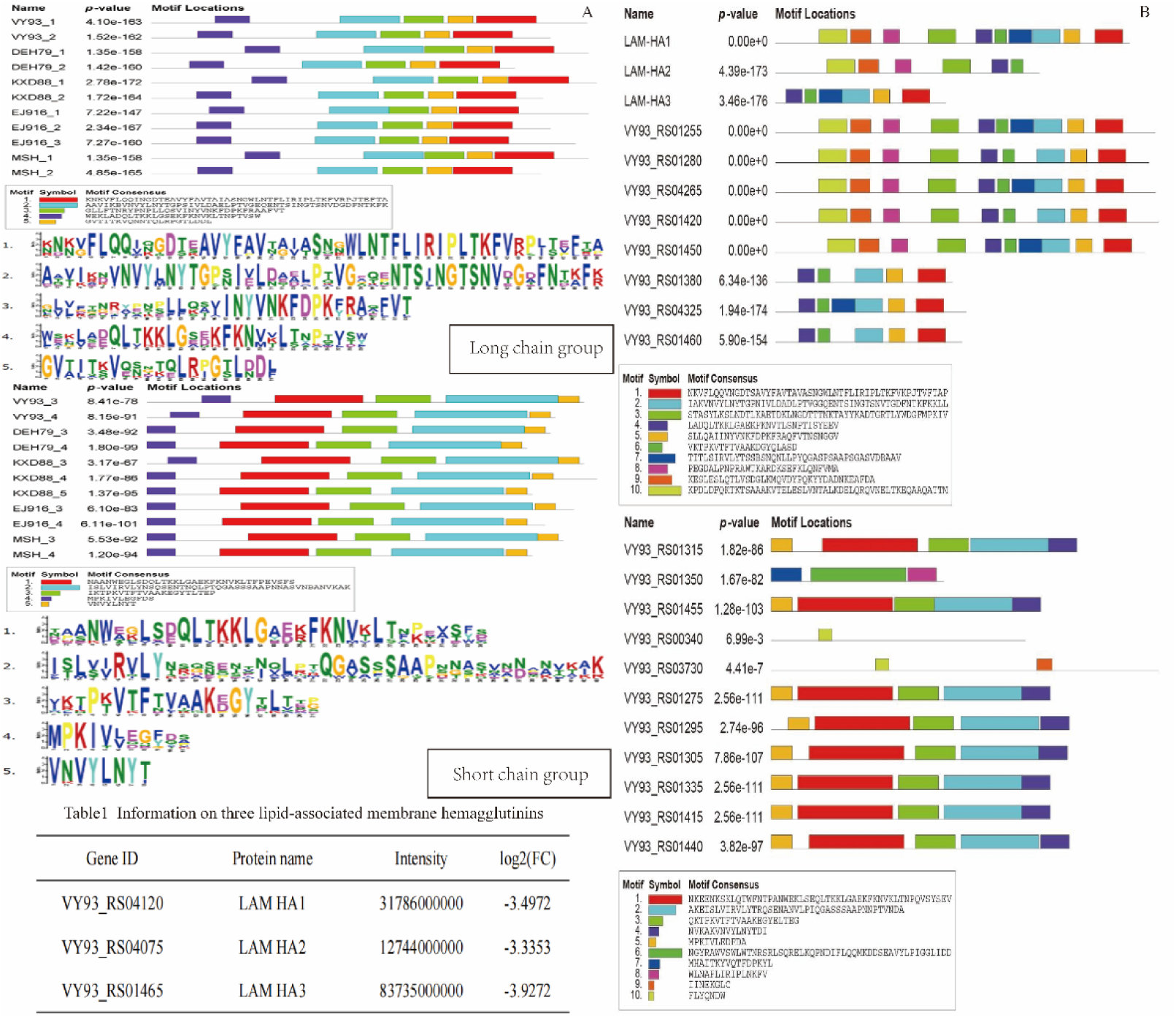
Visual analysis of the conserved structure of lipid-associated membrane hemagglutinin. (A) Analysis of conserved domains of M. synoviae hemagglutinin family. (B) Analysis of conserved domains of M. synoviae WVU1853 hemagglutinin family. The three hemagglutinins were identified as lipid-associated membrane hemagglutinins and were down-regulated when M. synoviae WVU1853 strain infected host cells.

### Eighteen host proteins interact with LAM HA screened by yeast two-hybrid

The yeast cDNA libraries were successfully constructed with host cells induced by *M. synoviae* with higher recombination rate (Fig 2B). The capacity of nuclear system secondary library was 1.44 × 10^7^ CFU (Fig 2C). The pGBKT7-LAM HA transformed Y2HGold and cDNA were screened in 3 rounds of deletion medium to obtain blue-positive clones, which were verified in line 2 rotations. The results showed that the transformed strains were able to grow on SD/-Trp/-Leu, SD/-Trp/-Leu/-His and SD/-Trp/-Leu/-His/-Ade plates and the colonies were blue in color on SD/-Trp/-Leu/-His/-Ade/X-α-Gal/3-AT plates in accordance with the results of the positive control (Fig 2D). The gene sequences of the positive bacteria were identified by sequencing (Fig 2E) and analyzed against the data in the NCBI database, and 18 proteins were obtained, as shown in Table 1 (Fig 2F).

**Fig 2.**
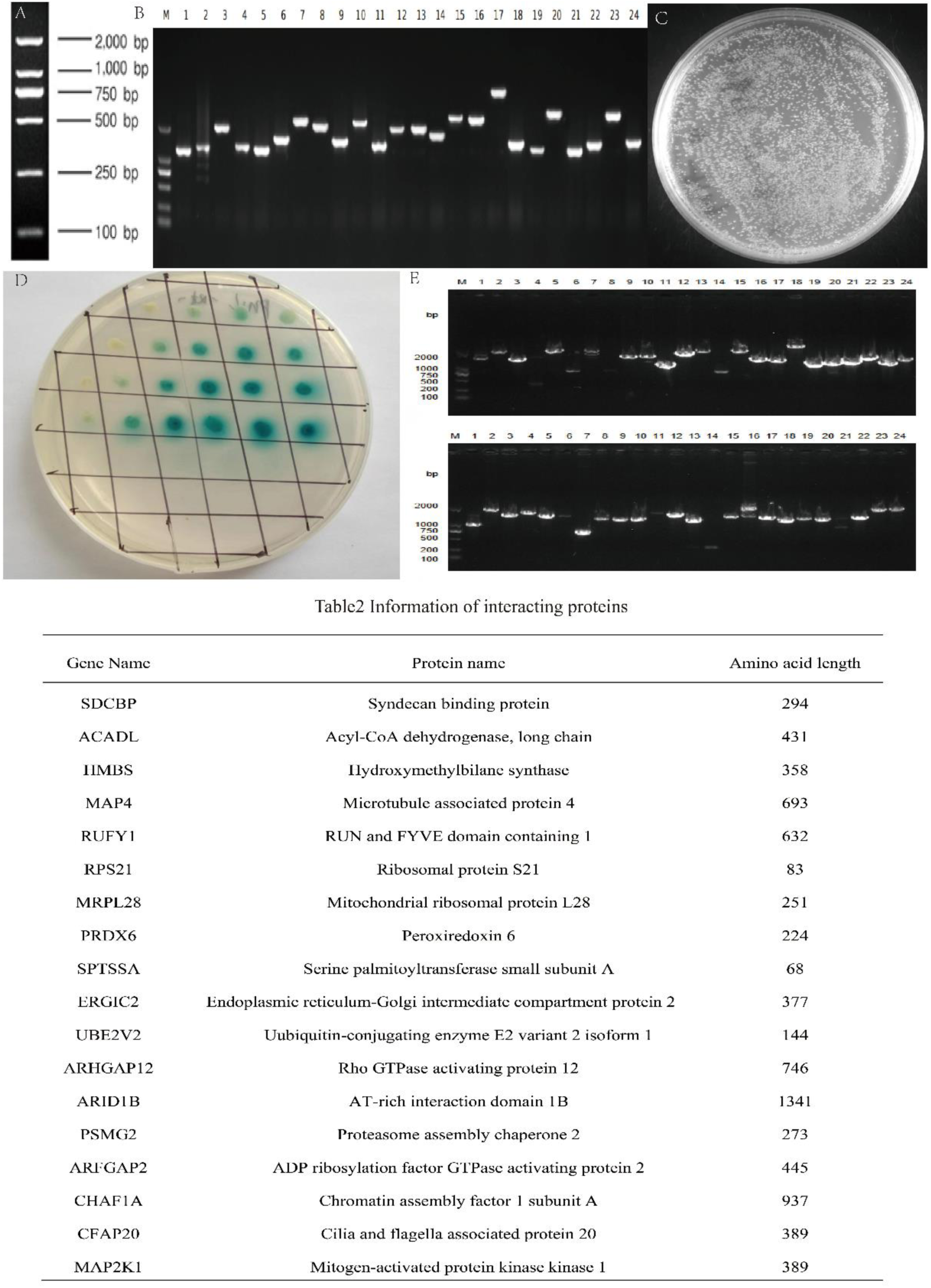
Yeast two-hybrid screening of host proteins interacting with lipid-associated membrane hemagglutinin. (A, B) The size of the inserted fragment of the constructed library was amplified by PCR, and the recombination rate was calculated by the number of colonies larger than 1000 bp accounting for the total number of colonies. (C) Identification of library insertion fragment size and recombination rate. (D) The size of the inserted fragment of the positive clone plasmid was amplified by PCR. The results of sequencing were compared and analyzed, and the information was listed in the table. (E) The plasmid with suitable sequencing was introduced into yeast containing pGBKT7-LAM HA for point-to-point verification.

### S83, R85, Y88, N124, and K192 are key amino acid residues mediating LAM Agglutination of erythrocytes

N15, S83 and Y88 appeared three times when forming complexes to form hydrogen bonds, and R85, N124 and K192 appeared four times, which were more frequent amino acid residues (Fig 3A). Successfully deleted five amino acids S83, Y88, R85, N124, and K192 and performed prokaryotic expression and purification, with a size of approximately 60 kD (Figure 3B). At pH 7.0 ∼ 7.5, LAM HA could induce chicken red blood cell agglutination, while ΔS83-R85-Y88-N124-K192-LAM HA did not induce chicken red blood cell agglutination (Fig 3C). When the pH was between 6.0 and 6.5, the agglutination titer of LAM HA was 1:2, and the agglutination titer of △S83-R85-Y88-N124-K192 LAM HA was 1:1; The agglutination titers of LAM HA and △S83-R85-Y88-N124-K192 LAM HA were both 1:1 when the pH was between 5 and 5.5 (Fig 3D).

**Fig 3.**
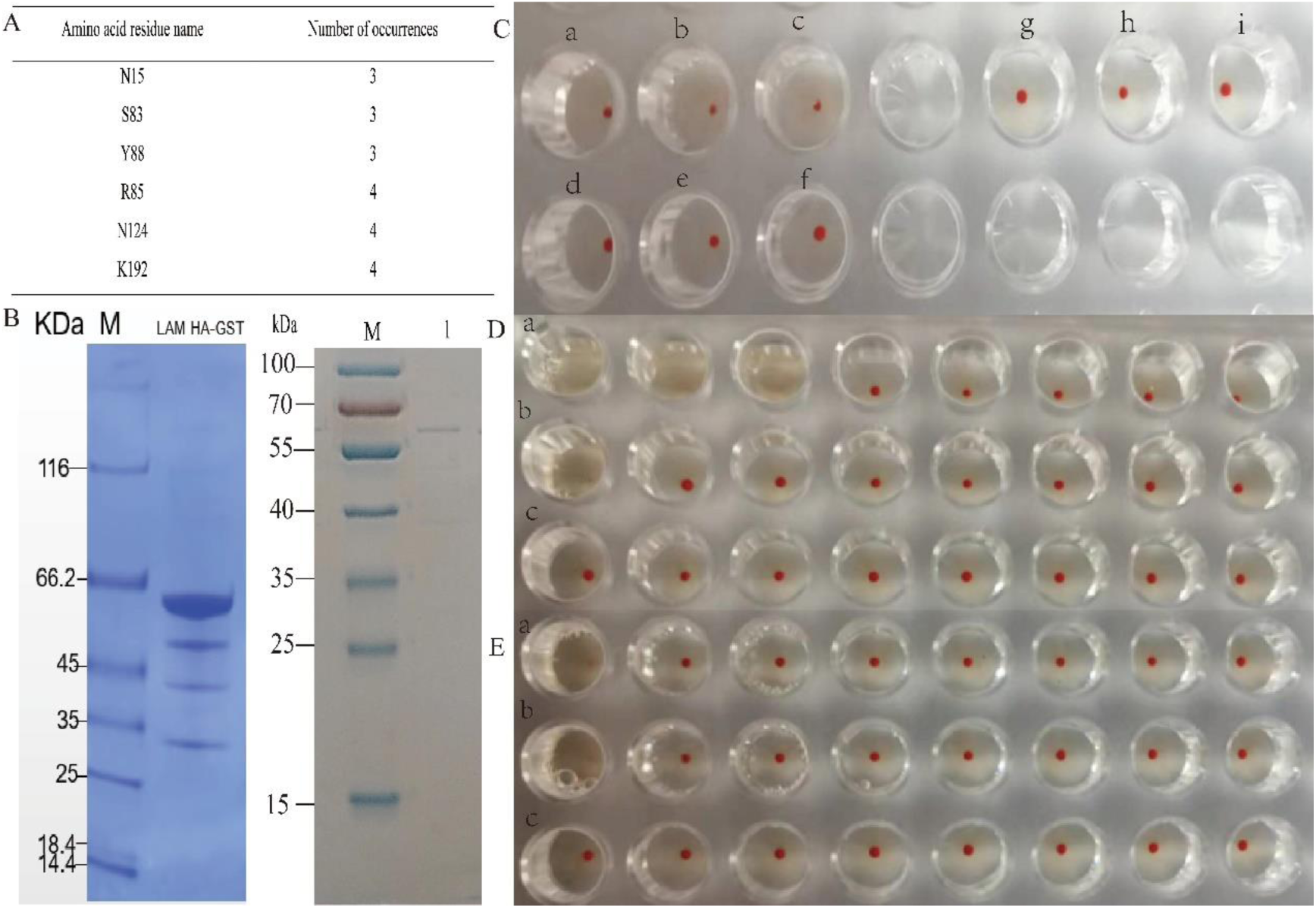
LAM HA is involved in the identification of key amino acid residues in agglutinating red blood cells. (A) LAM HA amino acid residues were involved in the statistics of the number of complex formation. (B) The deletion expression and purification of amino acid residues with high occurrence frequency of LAM HA. (C) Agglutination erythrocyte of LAM HA and △S83-R85-Y88-N124-K192-LAM HA at pH between 7.0 and 7.5. a, b and c are LAM HA coagulation tests, d, e and f are △S83-R85-Y88-N124-K192-LAM HA blood coagulation tests, and g, h and i are GST blood coagulation tests. (D) Agglutination erythrocyte of LAM HA and △S83-R85-Y88-N124-K192-LAM HA at pH between 6.0 and 6.5. Lines a, b and c were used to determine the hemagglutination titer of LAM HA, △S83-R85-Y88-N124-K192-LAM HA and GST, respectively. (E) Agglutination erythrocyte of LAM HA and △S83-R85-Y88-N124-K192-LAM HA at pH between 5.0 and 5.5. Lines a, b and c were used to determine the hemagglutination titer of LAM HA, △ S83-R85-Y88-N124-K192-LAM HA and GST, respectively.

### Analysis of the effect of pH on protein secondary structure

The results of the secondary structure prediction indicated that sialoreceptor binding motif PKVTFTVAAKDG mainly adopts a β-sheet structure.The residues S83, R85, Y88, and N124 are situated within the β-sheet region, while residue K192 is located within the α-helix region. Upon the deletion of residues S83, R85, Y88, N124, and K192, there was a decrease in the α-helix content, accompanied by an increase in the β-sheet and random coil content (Fig 4A, B). At a pH range of 6.0 to 6.5, spectroscopic analysis revealed a modest reduction in α-helix content and a slight increase in β-turn structures. Additionally, there was a decrease in β-sheet peptide formations (Fig 4 C, D).

**Fig 4.**
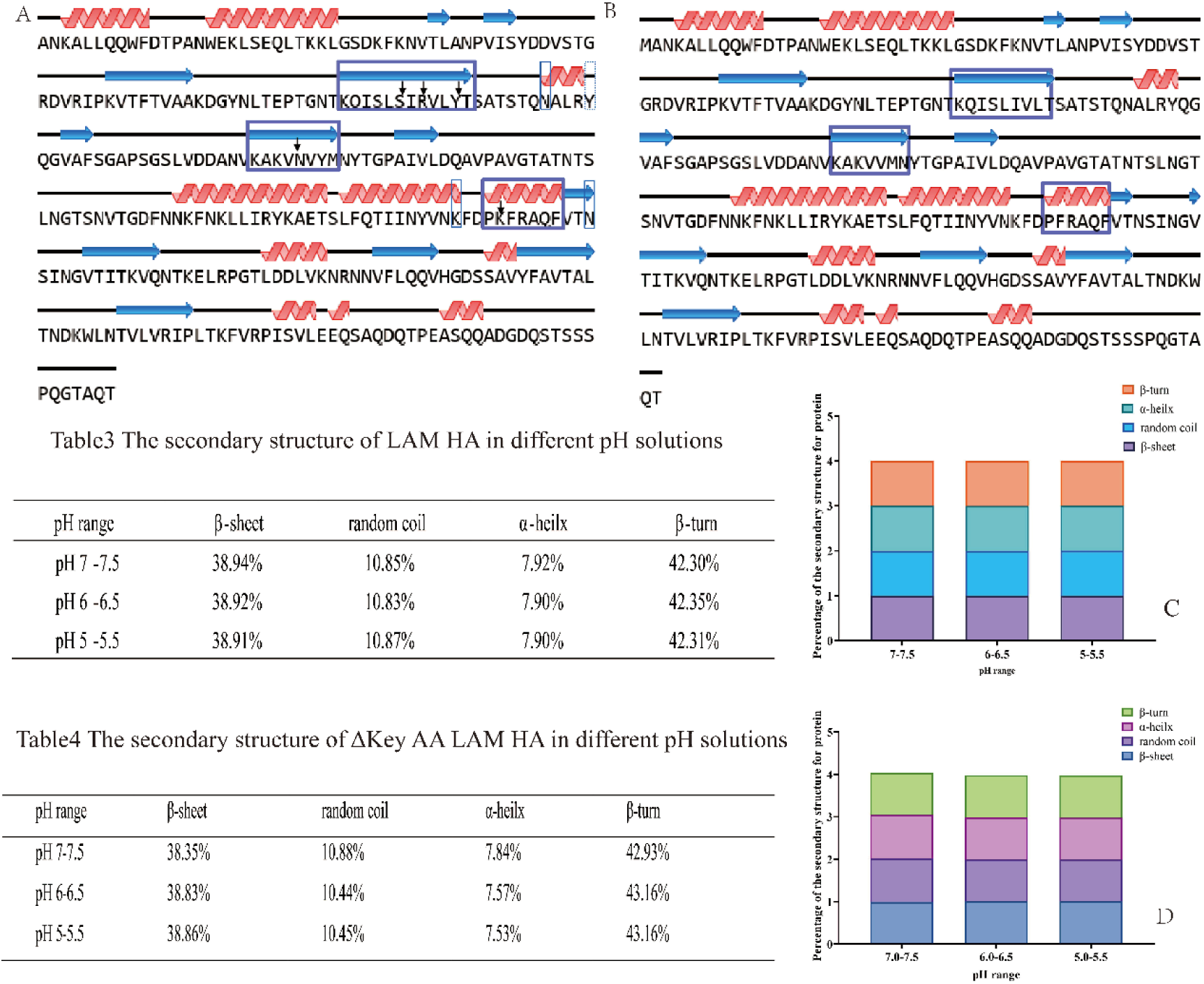
The effect of pH on the secondary structure of LAM HA and △S83-R85-Y88-N124-K192-LAM HA. (A) The secondary structure prediction of LAM HA. The arrow indicates the missing amino acid residue, and the purple box indicates the domain where the missing amino acid is located. The blue solid frame is an extra secondary structure, and the dotted frame indicates a secondary structure less than △S83-R85-Y88-N124-K192-LAM HA. (B) The secondary structure prediction of △ S83-R85-Y88-N124-K192-LAM HA. (C) The β-turn, α-helix, β-sheet and random coil proportion of LAM HA. (D) The β-turn, α-helix, β-sheet and random coil proportion of △S83-R85-Y88-N124-K192-LAM HA.

### Molecular dynamics analysis of differences in LAM HA and △ S83-R85-Y88-N124-K192-LAM HA structural stability

PyMOL analysis revealed that the RMSD values exceeded 3Å, indicating significant structural differences between the proteins before and after gene deletion (Fig 5A). Analysis of RMSD and RMSF across the trajectories revealed that under pH 5.0 conditions, the RMSD values of LAM HA were generally elevated, with a distinct canyon-shaped minimum observed at 2000 ps. Meanwhile, RMSF analysis revealed a significant peak in the region of residue 67-74 (sequence DGYNLTEPTG), whereas no such phenomenon was observed under the optimal pH 6.0 condition (Fig 5B). Under the same pH conditions, the RMSD values of the mutant △S83-R85-Y88-N124-K192 LAM HA were generally higher than those of the wild-type, with greater fluctuations observed at pH 5.0 and pH 6.0. At pH 7.0, however, its RMSD values showed little variation, although the minimum still occurred at the same time point. The RMSF values of the mutant were overall higher than those of the wild-type, particularly at pH 5.0 and pH 6.0. Specifically, the mutant exhibited significant peaks in the amino acid regions at positions 67-74, 24–32 (sequence TKKLGSDKF), and 141–150 (sequence GTATNTSLNGT). Furthermore, under pH 5.0 conditions, the mutant also exhibited a significant peak in the amino acid region spanning residues 108–116 (sequence PSGSLVDDANV) (Fig 5C).

**Fig 5.**
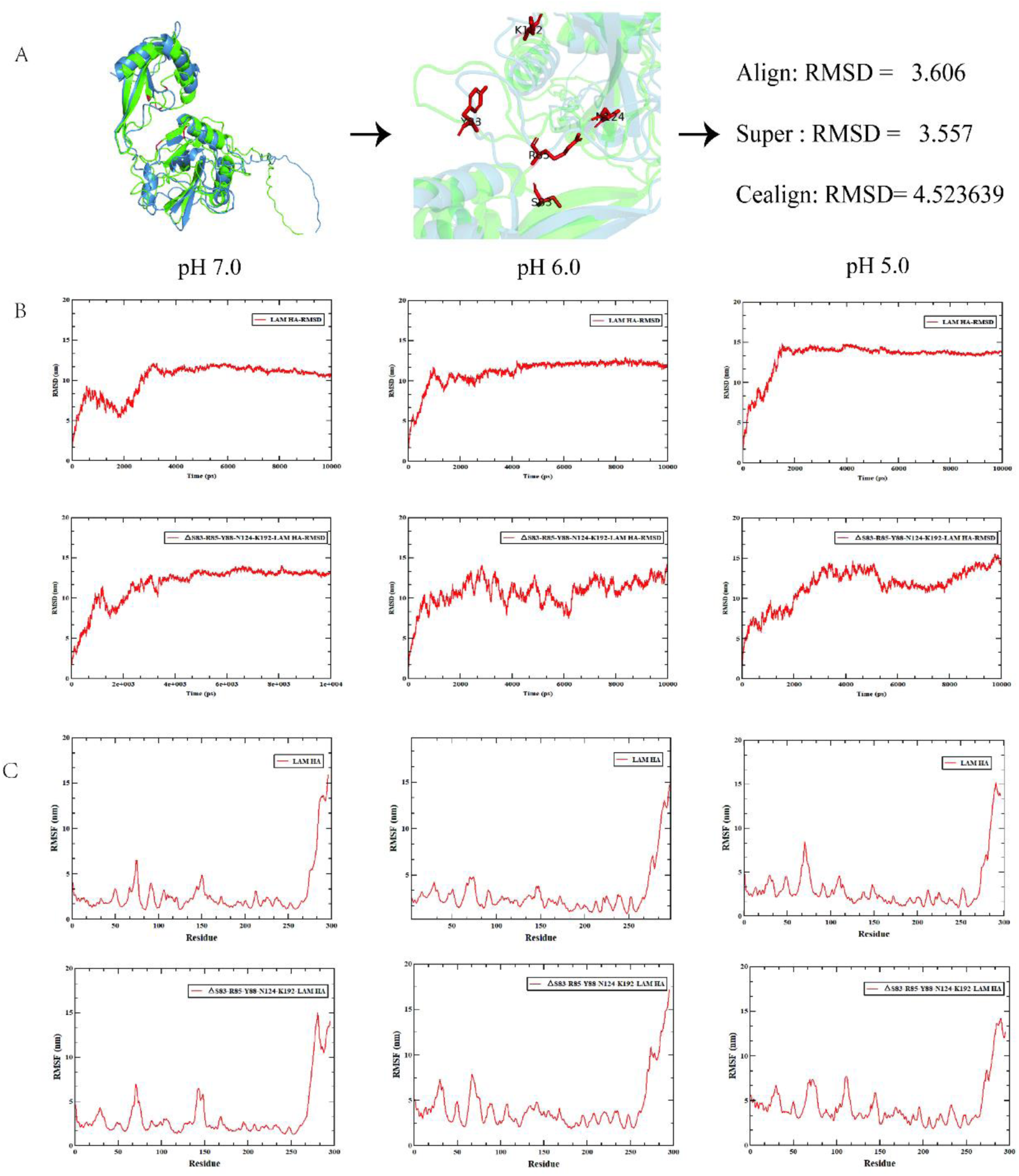
The stability prediction of LAM HA and △S83-R85-Y88-N124-K192-LAM HA. (A) RMSD assessment on LAM HA and △S83-R85-Y88-N124-K192-LAM HA. (B) RMSD assessment on LAM HA and △S83-R85-Y88-N124-K192-LAM HA under different pH conditions. (C) RMSF assessment on LAM HA and △S83-R85-Y88-N124-K192-LAM HA under different pH conditions.

## Discussion

High-throughput proteomic approaches, particularly the yeast two-hybrid (Y2H) system, have greatly advanced the mapping of protein–protein interaction networks due to their high sensitivity in detecting even weak or transient interactions ^(16, 17)^. To obtain a comprehensive host cDNA library reflecting infection-responsive expression, we constructed a Y2H library using RNA extracted from host cells infected with *M. synoviae* at multiple time points. Screening with LAM HA as bait identified 18 host protein interactors, five of which have known roles in pathogenic infection processes. These interactions illustrate how *M. synoviae* may manipulate host cell pathways to facilitate invasion and immune evasion ^(18)^. The interaction between proteins plays a vital role in the process of performing various biological functions. In most cases, due to the lack of experimentally determined complex structures, these interactions must be modeled to understand their molecular basis ^(14)^.

Similar to viral hemagglutinins, *M. synoviae* LAM HA binds receptors on erythrocytes, promoting colonization and dissemination in host tissues ^(19, 20)^. Although a conserved sialic acid–binding motif has been proposed ^(21)^, the functional domains of *M. synoviae* hemagglutinin remain poorly defined. Our molecular docking analyses repeatedly highlighted six residues-N15, S83, R85, Y88, N124, and K192 as potential functional determinants. N15 lies within an intein-related domain possibly involved in post-translational cleavage ^(22)^. S83, R85, and Y88 are situated in β-sheet regions, which in other hemagglutinins mediate heme binding and affect erythrocyte function ^(23, 24)^. N124 and K192 reside in α-helical structures known to facilitate protein adhesion ^(25)^. Targeted deletion of S83, R85, Y88, N124, and K192 significantly reduced but did not abolish hemagglutination, confirming their importance while suggesting additional residues contribute to activity. Notably, LAM HA function was pH-dependent, with enhanced activity under acidic conditions (pH 5.0–6.5), reminiscent of viral HA fusion activation in endosomal compartments ^(26, 27)^. Molecular dynamics simulations revealed that while LAM HA remains structurally stable across pH gradients, the ΔS83-R85-Y88-N124-K192 mutant exhibits marked conformational instability under acidic conditions. This computational observation aligns with circular dichroism data showing decreased β-sheet content and increased β-turn formation at pH 6.0–6.5, indicative of local unfolding or structural distortion ^(28, 29)^. At neutral pH 7.0–7.5, the mutant protein-though structurally intact-completely lost hemagglutination activity, indicating that the deleted residues are essential for maintaining the receptor-binding competent conformation ^(30)^. Under mild acidic conditions (pH 6.0–6.5), partial activity was restored alongside increased structural flexibility. We propose that protonation of key residues (e.g., HIP, ASH) in this pH range may induce an alternative, suboptimal conformation that permits low-affinity binding, resulting in reduced agglutination titers ^(31)^.

The hemagglutination titer assay results showed that wild-type LAM HA exhibited the highest activity at pH 6.0 (1:2), decreased activity at pH 5.0 (1:1), and was completely inactive at pH 7.0. The mutant △S83-R85-Y88-N124-K192 LAM HA showed an activity of 1:1 at both pH 6.0 and pH 5.0, and was inactive at pH 7.0. Combined with molecular dynamics simulation analysis, the molecular mechanism underlying this pH-dependent activity was elucidated. Simulation results showed that under pH 5.0 conditions, the RMSD values of wild-type LAM HA were generally elevated, with a ‘canyon’-shaped minimum occurring at 2000 ps. Meanwhile, RMSF analysis revealed a significant peak in the region of residue 67-74, whereas no such phenomenon was observed at the optimal pH 6.0. Sequence analysis revealed that this region is directly adjacent to the upstream sialylated receptor binding site (sequence PKVTFTVAAKDG), with RMSF values beginning to rise from the proline at the C-terminus of this binding site and peaking in the YNLTE region, indicating that flexibility is transmitted from the pH-sensitive loop to the receptor binding site. The region contains acidic residues such as Asp and Glu, with pKa values of approximately 4.0–4.5. At pH 6.0, these residues are fully deprotonated and negatively charged, stabilizing the local conformation through electrostatic interactions. When the pH drops to 5.0, partial protonation of the acidic residues leads to a significant increase in the flexibility of this region, which in turn induces global conformational perturbations and results in decreased hemagglutination activity. This phenomenon is consistent with the mechanism identified in recent studies, in which protonation of Asp/Glu under acidic conditions triggers protein desolvation and conformational rearrangement ^(32)^. The mutant exhibited generally elevated RMSD values with greater fluctuations compared to the wild-type at both pH 5.0 and pH 6.0, and overall higher RMSF values. Significant peaks were observed in the regions of residues 67-74, 24-32, 141-150, and 108-116 (at pH 5.0), indicating that the deleted S83-R85-Y88-N124-K192 region, particularly Y88, serves as a structural stability hub. This hub is spatially adjacent to the pH-sensitive loop, and its integrity is crucial for maintaining the overall conformational stability. Studies have shown that the pKa values of titratable residues within a protein are closely related to their local microenvironment, and the coupling between protonation state and conformational equilibrium represents a key mechanism for pH sensing and regulation ^(33, 34)^. Y88 may form aromatic interactions with the tyrosine in the YNLTE region, jointly maintaining the stability of this flexible region. Upon deletion of this region, even under the optimal pH 6.0 condition, the flexible loop at position 67-74 cannot be effectively stabilized, leading to a reduction in activity from 1:2 to 1:1. At pH 5.0, since the loop itself is already in a highly flexible state, the deletion has little effect on activity. At pH 7.0, both the wild-type and the mutant exist in inactive conformations. In summary, the pH-dependent hemagglutination activity of LAM HA is co-regulated by the pH-sensitive loop at position 67-74 and the S83-R85-Y88-N124-K192 structural stability hub, revealing a sophisticated regulatory mechanism of ‘local protonation-flexibility transmission-functional site perturbation’.

In conclusion, *M. synoviae* LAM HA functions as a key adhesion and hemagglutination protein, with its activity being critically dependent on specific residues (S83, R85, Y88, N124, K192) and pH conditions. These residues form a structural stability hub that, together with a pH-sensitive loop (residue 75), maintains an active conformation optimal at pH 6.0. Acidic pH shifts or deletion of these hub residues disrupts electrostatic interactions and local folding, leading to reduced or lost hemagglutination. These findings not only deepen our understanding of the molecular mechanism underlying LAM HA-mediated pathogenesis but also provide a theoretical basis for developing targeted interventions, vaccines, and diagnostic tools against *M. synoviae* infections.

## MATERIALS AND METHODS

### Bacterial strains and plasmids

Yeast two-hybrid library was constructed using pGADT7 plasmid and Y187 strain. The pGBKT7 plasmid and Y2HGold yeast strain were used to screen the library. PGEX-4t-1 plasmid used for prokaryotic expression, all of them were provided kindly by Clinical Veterinary Laboratory of Ningxia University. The target gene in the bait plasmid was synthesized by Shanghai Sangon and connected to the pGBKT7 vector.

### Hemagglutinin family of *M. synoviae* domain analysis

According to the hemagglutinin family related information integrated in the genome re-annotation ^(12)^ and lipid-related membrane protein identification experiments. The hemagglutinin sequences of 13 *M. synoviae* strains were extracted from the genome data published by NCBI (https://www.ncbi.nlm.nih.gov/datasets/genome/ (accessed on 29 September 2024)). Their conserved domains were analyzed with MEME (https://meme-suite.org/meme/ (accessed on 29 September 2024)).

### Bacterial two-hybrid library construction and screening

DF-1 and HD11 were infected with *M. synoviae* at 100 MOI, respectively. The culture medium was removed at 0 h, 4 h, 8 h, 12 h, and 24 h after infection, and PBS was washed three times. TRIzol was used to dissolve and collect cells. Then the RNA of each sample was extracted and determine the purity, the samples were mixed at a ratio of 1:1. The mixed samples were reverse transcribed into cDNA and a yeast two-hybrid library was constructed according to the Gateway method^(13)^.

For yeast two-hybrid screening, the LAM HA gene with *Bam*H I and *Xho* I restriction enzyme sites was amplified and cloned into the bait plasmid pGBKT7 to obtain pGBKT7-LAM HA. The pGBKT7-LAM HA and pGADT7 empty vector were co-transformed into Saccharomyces cerevisiae Y2HGold strain for self-activation detection. The bait plasmid pGBKT7-LAM HA was used to screen the candidate interacting proteins in the constructed chicken cell cDNA library. Plasmids of positive clones were screened from SD/-Trp-Leu-His-Ade defective plates containing 200 ng/ml Aureobasidin A (AbA). The positive plasmids were amplified by PCR using specific primers. The target fragment was sequenced and aligned to the chicken genome.

### Molecular docking of LAM HA with host proteins

SWISS-MODEL (https://swissmodel.expasy.org/interactive (accessed on 29 September 2024)) technology was used to generate a three-dimensional protein model through homology modeling for molecular docking of LAM HA and interacting host proteins. Cluspro (https://cluspro.org/help.php (accessed on 29 September 2024))^(14)^ was used to dock LAM HA with the successfully modeled interacting host protein. Simply, Beginning docking was used to open the PDB file of the target molecule, host protein was defined as the receptor molecule and LAM HA as the connecting molecule of molecular docking. PyMOL was used to visualize the interaction complex, amino acids that form hydrogen bonds within 3.5Å were counted in complex.

### Cloning and purification of recombinant proteins in *E. coli*

The LAM HA gene obtained from pGBKT7-LAM HA plasmid by *Bam*H I and *Xho* I digestion and cloned into pGEX-4T-1 to generate the plasmid pGEX-4T-LAM HA. △ Key AA LAM HA gene was synthesized by Sangon Biotech (Shanghai, China) and cloned into pGEX-4T-1 to generate the plasmid pGEX-4T-ΔLAM HA. The BL21/pGEX-4T-1, BL21/pGEX-4T-LAM HA and BL21/pGEX-4T-Δ LAM HA were inoculated in LB broth at 37°C at 220 rpm, until OD 600=0.7. The 0.5 mM final concentration isopropyl β-D-1-thioglutopropylglutaminoside (IPTG) was used for inducing protein expression. After 5 h of induction expression, at 16°C, 180 rpm, cells were collected by centrifugation at 10 000 rpm for 8 min at 4°C, resuspended and washed with PBS buffer three times. Then the resuspended cells with the Fisher Scientific™ Model 50 Sonic Dismembrator (Thermo Fisher Scientific Inc, USA) for 10 min. The supernatant centrifuge at 4°C, 12 000 rpm for 20 min to collect the solution. GST-tagged agarose magnetic bead method (Beyotime, China) was used to purify recombinant proteins. Recombinant proteins were desalted in dialysis tubing (Solaibao, China). The purified recombinant protein can be used for SDS-PAGE analysis.

### Hemagglutination test

LAM HA and △ Key AA LAM HA were diluted to 260 μg/mL. In a 96-well V-type micro-reaction plate, 0.025 mL PBS was added to each well from 2 to 8 wells in each row. The 1st wells were added with 0.025 mL LAM HA or △ Key AA-LAM HA, and repeatedly blown and sucked for 3-5 times to mix well. 0.025 mL of protein was extracted from the second hole and added to the third hole. After mixing, 0.025 mL of protein was extracted and added to the fourth hole, and then diluted to the seventh hole. 0.025 mL was extracted from the seventh hole and discarded. The eighth hole was the PBS control hole. Each well was added with 0.025 mL PBS, 0.025 mL 1% (volume fraction) chicken red blood cell suspension was added to each well. The reaction plate was mixed with reactants and allowed to stand at room temperature for 40 min. The results were determined when the red blood cells in the control hole were significantly button-like. The highest dilution of 100% agglutination was used to determine the hemagglutination titer. The pH of LAM HA and △ Key AA-LAM HA was adjusted to about 6.0 and 5.0, and the experiment was carried out according to the above steps to explore the effect of pH on erythrocyte agglutination.

### Fourier transform infrared (FTIR) spectroscopy

The secondary structure of the key amino acid residue sequence of LAM HA and △Key AA-LAM HA were visually analyzed by using Newpu online software (https://www.novopro.cn/tools/secondary-structure-prediction.html (accessed on 15 October 2024)). The proteins (5 mg/mL) for different pH were analysed using Nicolet™ iS50 FTIR spectroscopy (Thermo Fisher scientific., Waltham, USA). The dispersion medium was deionised phosphate buffer saline. Scanning was performed using a scanning infrared spectrometer at points between 400 and 4000 cm^−1^. Subsequently, data between 1600 and 1700 cm^−1^ were extracted for a Gaussian fit using Peakfit 4.12 software (Systat Software Inc., California, USA). The contents of the α-helix, β-sheet, β-turn and random coil were then calculated and visualized by GraphPad Prism (Version 10.2.0).

### Conformational analysis of LAM HA and △ Key AA-LAM HA at different pH

According to the amino acid sequence of the proteins, SWISS-MODEL was used to construct a three-level model and visualization by Pymol. The structures of LAM HA and △Key AA-LAM HA in different pH were analyzed by Amber software (Version Amber 25), The specific steps are as follows, in the obtained PDB file of each protein, HIS was changed to HID and named LAM HA pH 7.0 and △S83-R85-Y88-N124-K192 LAM HA pH 7.0, respectively. HIS was changed to HIP and ASP was changed to ASH, and these were named LAM HA pH 6.0 and △S83-R85-Y88-N124-K192 LAM HA pH 6.0, respectively. HIS was changed to HIP, GLU was changed to GLH, and ASP was changed to ASH, and these were named LAM HA_pH 5.0 and △S83-R85-Y88-N124-K192 LAM HA pH 5.0, respectively. After loading the leaprc.protein.ff19SB force field and balancing the charges, the topology and trajectory files for each protein were successfully generated. Then the root mean square deviation (RMSD) and root mean square fluctuation (RMSF) of each structure were analyzed by CPPTRAJ Software package ^(15)^. Subsequently, qtgrace software (qtgrace_v026_Win7) was used for visualization.

## ACKNOWLEDGMENTS

We thank Data Institute Server. This work was also supported by the Department of Science and Technology of Ningxia Hui Autonomous Region and Ningxia Xiaoming Agriculture and Animal Husbandry Co., Ltd.

This work was supported by the [Postdoctoral research start-up project] to D.D.S., [Postdoctoral Science Foundation Project] to D.D.S., [Ningxia Hui Autonomous Region Science and Technology Innovation Team Building Project], funding number [2022BSB03107] to J.D.L.

The funders had no role in the study design, data analysis, data interpretation, and the writing of this manuscript. All authors had full access to the study data and took responsibility for submitting it for publication.

D.D.S., S.H.H., and J.D.L. conceived and designed the study. D.D.S. performed the experiments and analyzed the data. L.G., X.H.C., X.X.H., and Q.W. participated in the implementation of the study. D.D.S., F.Q.W., and Y.S. wrote the manuscript and produced the figures. S.J.B., S.H.H., L.G., X.H.C., and J.D.L. critically revised and edited the manuscript. D.D.S., and J.D.L. secured the funds. S.J.B., S.H.H., and L.G. supervised the project. All authors reviewed and approved the final version of the manuscript.

## Contributor Information

Jing-Duan Li.

Sheng-Hu He

## DATA AVAILABILITY

Data availability statement is not applicable to this study because no high-throughput omics datasets (genomics, transcriptomics, proteomics, etc.) were generated or analyzed.

## ETHICS APPROVAL

This study was approved by the Animal Research Ethics Committee of Gansu Agricultural University (Permit No. 2024-043).

